# The structural connectome constrains fast brain dynamics

**DOI:** 10.1101/2020.11.25.393017

**Authors:** P Sorrentino, C Seguin, R Rucco, M Liparoti, E Troisi Lopez, S Bonavita, M Quarantelli, G Sorrentino, V Jirsa, A Zalesky

**Affiliations:** Aix-Marseille University, Inserm, INS, Institut de Neurosciences des Systèmes, Marseille, France; Institute of Applied Sciences and Intelligent Systems, National Research Council, Pozzuoli, Italy; University of Melbourne, Melbourne, Australia; Department of Motor Sciences and Wellness, Parthenope University of Naples, Naples, Italy; Institute for Diagnosis and Cure Hermitage Capodimonte, Naples, Italy; University of Campania Luigi Vanvitelli. Caserta, Italy; Biostructure and Bioimaging Institute, National Research Council, Naples, Italy

**Keywords:** brain dynamics, brain networks, magnetoencephalography, neuronal avalanches, structural connectome

## Abstract

Brain activity during rest displays complex, rapidly evolving patterns in space and time. Structural connections comprising the human connectome are hypothesized to impose constraints on the dynamics of this activity. Here, we use magnetoencephalography (MEG) to quantify the extent to which fast neural dynamics in the human brain are constrained by structural connections inferred from diffusion MRI tractography. We characterize the spatio-temporal unfolding of whole-brain activity at the millisecond scale from source-reconstructed MEG data, estimating the probability that any two brain regions will significantly deviate from baseline activity in consecutive time epochs. We find that the structural connectome relates to, and likely affects, the rapid spreading of neuronal avalanches, evidenced by a significant association between these transition probabilities and structural connectivity strengths (r=0.37, p<0.0001). This finding opens new avenues to study the relationship between brain structure and neural dynamics.

## Introduction

The structural scaffolding of the human connectome (1) constrains the unfolding of large-scale coordinated neural activity towards a restricted *functional repertoire* (2). While functional magnetic resonance imaging (fMRI) can elucidate this phenomenon at relatively slow timescales (3–5), brain activity shows rich dynamic behaviour across multiple timescales, with faster activity nested within slower scales. Here, in healthy young adults, we exploit the high temporal resolution of resting-state magnetoencephalography (MEG) data to study the spatial spread of perturbations of local activations representative of neuronal avalanches. We aim to establish whether the structural connectome constrains the spread of avalanches among regions (6, 7). We find that avalanche spread is significantly more likely between pairs of grey matter regions that are structurally connected, as inferred from diffusion MRI tractography. This result provides cross-modal empirical evidence suggesting that connectome topology constrains fast-scale transmission of neural information, linking brain structure to brain dynamics.

## Results

Structural connectomes were mapped for 58 healthy adults (26 females, mean age ± SD: 30.72 ± 11.58) using diffusion MRI tractography and regions defined based on the Automated Anatomical Labeling (AAL) and the Desikan-Killiany-Tourville (DKT) atlases. Interregional streamline counts derived from whole-brain deterministic tractography quantified the strength of structural connectivity between pairs of regions. Streamline counts were normalized by regional volume. Group-level connectomes were computed by averaging connectivity matrices across participants.

MEG signals were pre-processed and source reconstructed for both the AAL and DKT atlases. All analyses were conducted on source-reconstructed signal amplitudes. Each signal amplitude was z-scored and binarized such that, at any time point, a z-score exceeding a given threshold was set to 1 (active); all other timepoints were set to 0 (inactive). An avalanche was defined as starting when any region exceeded this threshold, and finished when no region was active. An avalanche-specific transition matrix (TM) was calculated, where element (*i, j*) represented the probability that region *j* was active at time *t+*◻, given that region *i* was active at time *t*, where ◻∼3ms. The TMs were averaged per participant, and then per group, and finally symmetrized. Fig.1 provides an overview of the pipeline.

**Figure 1.**
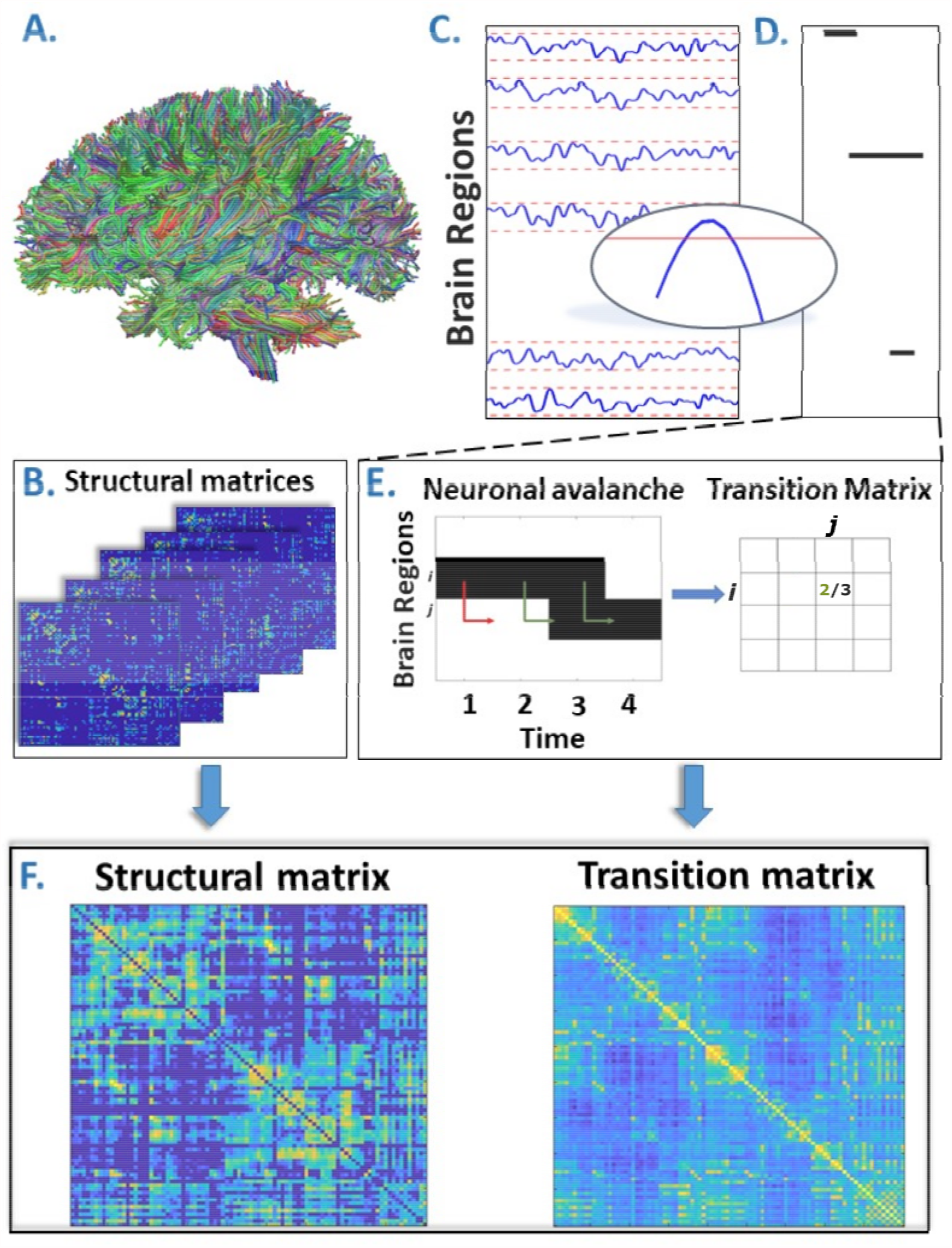
A. Rendering of streamlines reconstructed using diffusion MRI and tractography for an individual. B. Structural connectivity matrix. Row/columns represent regions comprising a brain atlas. Matrix entries store the number of streamlines interconnecting each pair of regions. C. Source-reconstructed MEG series. Each blue line represents the z-scored activity of a region, and the red lines denote the threshold (z-score= ± 3). The inset represents a magnified version of a time-series exceeding the threshold. D. Raster plot of an avalanche. For each region, the moments in time when the activity is above threshold are represented in black, while the other moments are indicated in white. The particular avalanche that is represented involved three regions. E. Estimation of the transition matrix of a toy avalanche. Region *i* is active three times during the avalanche. In two instances, denoted by the green arrows, region *j* was active after region *i*. In one instance, denoted by the red arrow, region *i* is active but region *j* does not activate at the following time step. This situation would result, in the transition matrix, as a 2/3 probability. F. Average structural matrix and average transition matrix (Log scale).

We found striking evidence of an association between avalanche transition probabilities and structural connectivity strengths (Fig. 2), suggesting that regional propagation of fast-scale neural avalanches is partly shaped by the axonal fibers forming the structural connectome (r=0.40, p<0.0001). Specifically, the association was evident for different activation thresholds and both the AAL and DKT connectomes (AAL atlas: for threshold z=2.5, r=0.41; for threshold z=3.0, r=0.40; for threshold z=3.5, r=0.39; DKT atlas: for threshold z=2.5, r=0.38; for threshold z=3.0, r=0.37; for threshold z=3.5, r=0.35; in all cases, p <0.0001), as well as for individual- and group-level connectomes, although associations were stronger for group-level analyses (see Fig. 2, panel A).

**Figure 2.**
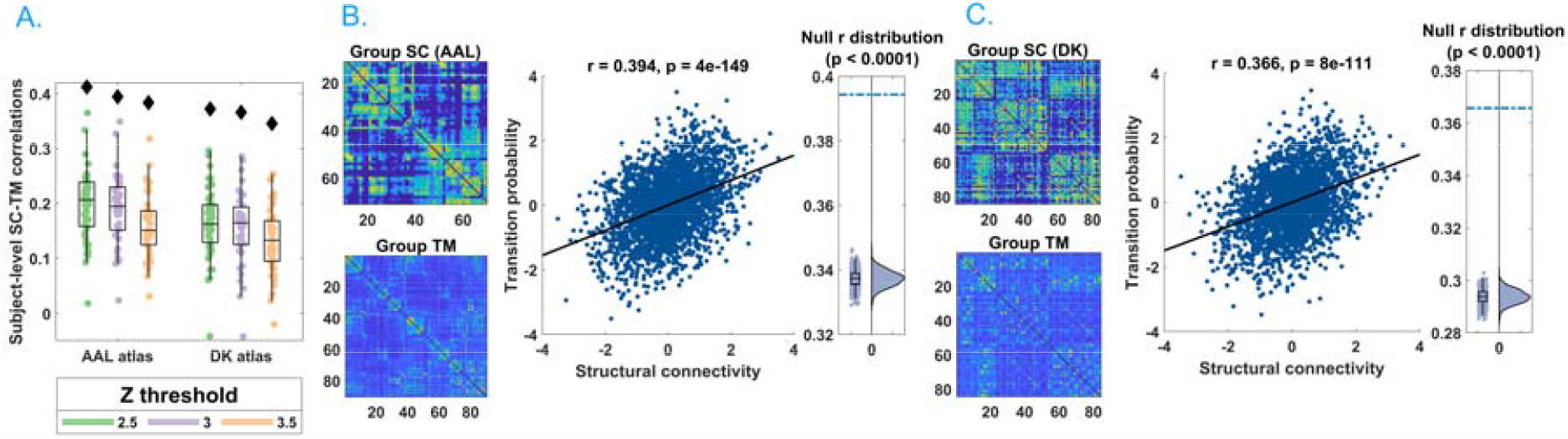
A. Distribution of the r’s of the Spearman’s correlation between the subject-specific transition matrices and structural connectomes. The black diamond represent the r’s of the group-averaged matrices. On the left, the results for the AAL atlas, on the right, the results for the DKT atlas. Green, purple and orange dots represent results obtained with a z-score threshold of 2.5, 3 and 3.5, respectively. **B and C**. Data referring to the AAL atlas in B, to the DKT atlas in C. On the top-left, the average structural matrix, on the bottom left, the average transition matrix. The scatterplot shows the correlation between the values of the structural edges and the transition probabilities for the corresponding edge. The black line represents the best fit line in the least-square sense. On the right, the distribution shows the r’s derived from the null distribution. The dotted blue line represents the observed r. Please note that, for visualization purposes, the connectivity weights and the transition probabilities were resampled to normal distributions. Figure 2-figure supplement 1 shows the comparison between the structural connectome and the transition matrix computed by taking into account longer delays. In the Supplementary file 1, we report a table with an overview of the results of the frequency-specific analysis.

We also investigated this phenomenon within specific frequency bands. Associations were evident in all the classical frequency bands: delta (0.5 – 4 Hz; r=0.39), theta (4 – 8 Hz; r=0.29), alpha (8 – 13 Hz; r=0.32), beta (13 – 30 Hz; r=0.32), and gamma (30 – 48 Hz; r=0.32), with p<0.0001 for all bands (see Supplementary File 1). Supplementary analyses suggested that these results could not be attributable to volume conduction confounds (see Methods; Field spread analysis).
Next, we sought to test whether the associations were weaker for randomized transition matrices computed after randomizing the times of each avalanche while keeping the spatial structure unchanged. Randomized transition matrices resulted in markedly weaker associations with structural connectivity, compared to the actual transition matrices (AAL atlas, z-score=3: mean r = 0.26, observed r= 0.40, p<0.001). Note that the mean correlation coefficient was greater than zeros for the randomized data because the randomization process preserved basic spatial attributes in the data. We also found that the findings remained significant after excluding subcortical regions (with lower signal-to-noise ratios). Finally, we replicated these findings for a group-level connectome derived using diffusion MRI acquired from 200 healthy adults in the Human Connectome Project (r=0.11, p<0.001, z-score=3; see Methods). Our results were thus robust to multiple connectome mapping pipelines and parcellation atlases, significant for both group-averaged and individual connectomes, and could not be explained by chance transitions and/or volume conduction effects. Collectively, these results suggest that connectome organization significantly shapes the propagation of neural activity.

## Discussion

Our results provide new insight into the propagation of fast-evolving brain activity in the human connectome. We show that the spatial unfolding of neural dynamics at the millisecond scale relates to the network of large-scale axonal projections comprising the connectome, likely constraining the exploration of the brain’s putative functional repertoire. The short time scale of several milliseconds biases the constraint to direct connections, which is the focus of this paper. Longer delays may impose constraints upon larger-scale motifs of the network and further characterize the sub-spaces, in which brain dynamics unfold.

Previous functional MRI studies provide evidence of coupling between structural connectivity and slow activations (3, 8, 9). However, intrinsic neural dynamics evolve quickly and are nested within slow activity (10). Our findings suggest that long-term structure-function coupling occurs against a backdrop of faster fluctuations, which are also constrained by the connectome and may enable individuals to rapidly respond to changing environments and new cognitive demands (11).

Consistent with our findings, two recent M/EEG studies showed that functional connectivity, as estimated using amplitude-envelope coupling (AEC), relates to structural connectivity (12, 13). However, in contrast to AEC, we conducted time-resolved analyses, characterizing avalanche dynamics at high temporal resolution. Further work is needed to determine the extent to which structure-function coupling is dynamic. To this regard, our results suggest that coupling is strongest during avalanche events, consistent with established theories (14). Finally, our results might explain how the large-scale activity unfolding in time might lead to the previous observation that average resting-state functional connectivity displays topological features that mirror those of the structural connectome (15). Our proposed framework links the large-scale spreading of aperiodic, locally generated perturbations to the structural connectome, and might be further exploited to investigate polysynaptic models of network communication, which aim to describe patterns of signalling between anatomically unconnected regions (16, 17). In fact, our results show that transitions of activations are observed across regions that do not appear to be directly linked in the structural connectome. This provides evidence for polysynaptic communication.

Neuronal avalanches have been previously observed in MEG data (7), and their statistical properties, such as a size distribution that obeys a power-law with an exponent of -3/2, reported. These features are compatible with those that would be predicted starting from a process operating at criticality with a branching ratio equal to one. While beyond the scope of this paper, our framework might contribute to elucidating the role of the structural scaffolding (and its topological properties) to the emergence of the observed large-scale, scale-free critical dynamics. In turn, this might be exploited to predict the effects of structural lesions on behaviour and/or clinical phenotypes.

While our findings were replicated across multiple frequency bands, structural connectivity can potentially impose frequency-dependent constraints on avalanche spread. Future work should investigate frequency-specific data to understand what leads to the emergence of avalanches and, most importantly, to the specific spatio-temporal patterns of recruited regions that defines individual (or at least groups of) avalanches in each specific frequency-band.
For the present application, we reconstructed the structural connectome using a deterministic tractography algorithm. While probabilistic algorithms can provide advantages in some applications, they are prone to reconstruction of spurious connections (false positives), compared to deterministic methods, reducing connectome specificity (18, 19). We used deterministic tractography because previous functional MRI studies report that structure-functional coupling is greater for connectivity matrices inferred from deterministic tractography, compared to probabilistic methods (20). Nonetheless, additional studies are needed to clarify if and to what extent the present results are influenced by the structural connectome reconstruction method. While we replicated our findings using alternative datasets (i.e. HCP) and parcellations, further replication using alternative connectome mapping pipelines is warranted.

In conclusion, using MEG to study fast neuronal dynamics and diffusion MRI tractography to map connectomes, we found that the connectome significantly constrains the spatial spread of neuronal avalanches to axonal connections. Our results suggest that large-scale structure-function coupling is dynamic and peaks during avalanche events.

## Methods

### Participants

We recruited 58 young adults (male 32 / female 26, mean age ± SD was 30.72 ± 11.58) from the general community. All participants were right-handed and native Italian speakers. The inclusion criteria were: 1) no major internal, neurological or psychiatric illnesses; 2) no use of drugs or medication that could interfere with MEG/MRI signals. The study complied with the Declaration of Helsinki and was approved by the local Ethics Committee. All participants gave written informed consent.

### MRI acquisition

3D T1-weighted brain volumes were acquired at 1.5 Tesla (Signa, GE Healthcare) using a 3D Magnetization-Prepared Gradient-Echo BRAVO sequence (TR/TE/TI 8.2/3.1/450 ms, voxel 1 × 1 × 1 mm3, 50% partition overlap, 324 sagittal slices covering the whole brain), and diffusion MRI data for individual c connectome reconstruction were obtained using the following parameters: Echo-Planar Imaging, TR/TE 12,000/95.5 ms, voxel 0.94×0.94×2.5mm3, 32 diffusion-sensitizing directions, 5 B0 volumes). The MRI scan was performed after the MEG recording. Preprocessing of the diffusion MRI data was carried out using the software modules provided in the FMRIB Software Library (FSL, http://fsl.fmrib.ox.ac.uk/fsl). All diffusion MRI datasets were corrected for head movements and eddy currents distortions using the “eddy_correct” routine, rotating diffusion sensitizing gradient directions accordingly, and a brain mask was obtained from the B0 images using the Brain Extraction Tool routine. A diffusion-tensor model was fitted at each voxel, and streamlines were generated over the whole brain by deterministic tractography using Diffusion Toolkit (FACT propagation algorithm, angle threshold 45°, spline-filtered, masking by the FA maps thresholded at 0.2). For tractographic analysis, the ROIs of the AAL atlas and of a MNI space-defined volumetric version of the Desikan-Killiany-Tourville (DKT) ROI atlas were used, both masked by the GM tissue probability map available in SPM (thresholded at 0.2). To this end, for each participant, FA volumes were normalized to the MNI space using the FA template provided by FSL, using the spatial normalization routine available in SPM12, and the resulting normalization matrices were inverted and applied to the ROIs, to apply them onto each subject. The quality of the normalization was assessed visually. From each subject’s whole brain tractography and corresponding GM ROI set, the number of streamlines connecting each couple of GM ROIs and the corresponding mean tract length was calculated using an in-house software written in Interactive Data Language (IDL, Harris Geospatial Solutions, Inc., Broomfield, CO, USA).

Connectomes in the replication dataset were constructed using an alternative mapping pipeline and diffusion MRI data from the Human Connectome Project (HCP). Deterministic tractography was performed using MRtrix3 (21) under the following parameters: FACT algorithm, 5 million streamlines, 0.5 mm propagation step size, 400 mm maximum propagation length, and 0.1 FA threshold for the termination of streamlines (17). The number of streamlines connecting any couple of regions was normalized by the combined volume of the two regions. Structural matrices were constructed for 200 HCP participants using the AAL atlas and averaged to derive a group-level connectome.

### MEG pre-processing

MEG pre-processing and source reconstruction were performed as in (22). The MEG system was equipped with 163 magnetometers, and was developed by the National Research Council of Italy at the Institute of Applied Sciences and Intelligent Systems (ISASI). All technical details regarding the MEG device are reported in (23). In short, the MEG registration was divided in two eyes-closed segments of 3:30 minutes each. To identify the position of the head, four anatomical points and four position coils were digitized. Electrocardiogram (ECG) and electro-oculogram (EOG) signals were also recorded. The MEG signals, after an anti-aliasing filter, were acquired at 1024 Hz, then a fourth order Butterworth IIR band-pass filter in the 0.5-48 Hz band was applied. To remove environmental noise, measured by reference magnetometers, we used Principal Component Analysis. We adopted supervised Independent Component Analysis to clean the data from physiological artifacts, such as eye blinking (if present) and heart activity (generally one component). Noisy channels were identified and removed manually by an expert rater (136 ± 4 sensors were kept). 47 subjects were selected for further analysis.

### Source reconstruction

The time series of neuronal activity were reconstructed in 116 regions of interests (ROIs) based on the Automated Anatomical Labeling (AAL) atlas (24, 25); and in 84 regions of interest based on the Desikan-Killiany-Tourreville (DKT) atlas. To do this, we used the volume conduction model proposed by Nolte (26) applying the Linearly Constrained Minimum Variance (LCMV) beamformer algorithm (27) based on the native structural MRIs. Sources were reconstructed for the centroids of each ROI. Finally, we considered a total of 90 ROIs for the AAL atlas, since we have excluded 26 ROIs corresponding to the cerebellum because of their low reliability in MEG (28). All the preprocessing steps and the source reconstruction were made using the Fieldtrip toolbox (29).

### Neuronal avalanches and branching parameter

To study the dynamics of brain activity, we estimated “neuronal avalanches”. Firstly, the time series of each ROI was discretized calculating the z-score, then positive and negative excursions beyond a threshold were identified. The value of the threshold was set to 3 standard deviations (|*z*| = 3), but we tested the robustness of the results changing this threshold from 2.5 to 3.5. A neuronal avalanche begins when, in a sequence of contiguous time bins, at least one ROI is active (|*z*| >3), and ends when all ROIs are inactive (30, 31). The total number of active ROIs in an avalanche corresponds to its size.

These analyses require the time series to be binned. This is done to ensure that one is capturing critical dynamics, if present. To estimate the suitable time bin length, for each subject, for each neuronal avalanches and for each time bin duration, the branching parameter σ was estimated (32, 33). In fact, system operating at criticality typically display a branching ratio ∼1. The branching ratio is calculated as the geometrically averaged (over all the time bins) ratio of the number of events (activations) between the subsequent time bin (descendants) and that in the current time bin (ancestors) and then averaging it over all the avalanches (34). More specifically:

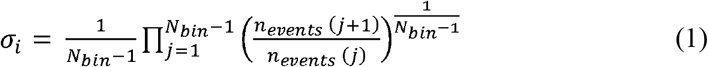

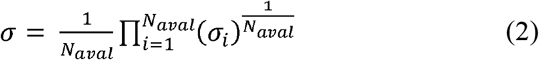

Where *σ*_*i*_ is the branching parameter of the i-th avalanche in the dataset, *N*_*bin*_ is the total amount of bins in the i-th avalanche, *n*_*events*_ (*j*) is the total number of events active in the j-th bin, *N*_*aval*_ is the total number of avalanche in the dataset. We tested bins from 1 to 5, and picked 3 for further analyses, given that the branching ratio was 1 for bin =3. However, results are unchanged for other bin durations, and the branching ratio remains equal to 1 or differences were minimal (range: 0.999 to 1.010 - data not shown). Bins of longer duration would violate the Nyquist criterion and were thus not considered. The results shown are derived when taking into accounts avalanches longer than 10 time bins. However, we repeated the analysis taking into account avalanches longer than 30 time bins, as well as taking all avalanches into account, and the results were unchanged.

### Transition matrices

The amplitude of each binned, z-scored source-reconstructed signal was binarized, such that, at any time bin, a z-score exceeding ± 3 was set to 1 (active); all other time bins were set to 0 (inactive). Alternative z-score thresholds (i.e. 2.5 and 3.5) were tested. An avalanche was defined as starting when any region is above threshold, and finishing when no region is active, as in (22). Avalanches shorter than 10 time bins (∼30 msec) were excluded. However, the analyses were repeated including only avalanches longer than 30 time bins (∼90 msec), to focus on rarer events (sizes of the neuronal avalanches have a fat-tailed distribution) that are highly unlikely to be noise, and including all avalanches, and the results were unchanged. An avalanche-specific transition matrix (TM) was calculated, where element (*i, j*) represented the probability that region *j* was active at time *t+*◻, given that region *i* was active at time *t*, where ◻∼3ms. The TMs were averaged per participant, and then per group, and finally symmetrized. The introduction of a time-lag makes it unlikely that our results can be explained trivially by volume conduction (i.e. the fact that multiple sources are detected simultaneously by multiple sensors, generating spurious zero-lags correlations in the recorded signals). For instance, for a binning of 3, as the avalanches proceed in time, the successive regions that are recruited do so after roughly 3 msecs (and 5 msecs for the binning of 5). Hence, activations occurring simultaneously do not contribute to the estimate of the transition matrix. See below for further analyses addressing the volume conduction issue. Finally, we explored transition matrices estimated using frequency-specific signals. To this end, we filtered the source-reconstructed signal in the classical frequency bands (delta, 0.5 – 4 Hz; theta 4 – 8 Hz; alpha 8 – 13 Hz; beta 13 – 30 Hz; gamma 30 – 48 Hz), before computing neuronal avalanches and the transition matrix, by applying a fourth-order Butterworth pass-band filter to the source-reconstructed data, before proceeding to the further analysis as previously described. The results remained significant in all the explored frequency bands. This analysis was carried out for the DKT atlas, binning = 3, z-score threshold = ± 3.

#### Field spread analysis

Volume conduction alone is an unlikely explanation of our results, given that simultaneous activations do not contribute to the transition matrix, due to the time lags introduced. To confirm that volume conduction effects were negligible, the transition matrices were re-computed using longer delays. In short, we identified the regions that were recruited in an avalanche after the first perturbation (i.e. the initial time-bin of an avalanche). Since we did not scroll through the avalanche in time, as previously described, we considered time delays as long as the avalanche itself, while minimizing the influence of short delays. This means that the avalanche-specific transition matrix is now binary, and the *ij*^*th*^ element is equal to 1 if region *i* started the avalanche (i.e. it was active at the first time-bin) and region *j* was recruited in the avalanche at any subsequent timepoint, and 0 otherwise. This alternative procedure for the estimation of the transition matrices was carried out for the AAL atlas, in the case of binning =3, z-score threshold = ± 3. In this case, a significant association remained between transition probabilities and structural connectivity (r=0.36; p<0.0001). Figure 2–figure supplement 1 provides further details.

To further rule out the possibility that field spread might introduce spurious correlations that might drive the relationship between the Transition Matrix and the structural connectivity matrix, we conducted further analyses involving surrogate data. We generated n white Gaussian processes, with n = 66, i.e. the number of cortical regions, and we smoothed them using a zero-phase polynomial filter. Then, we added 100 perturbations, where each perturbation was assigned to a randomly chosen regions and random time point, subject to the following constraints. Perturbations were separated by at least 200 samples (no overlap was allowed, i.e. the perturbations could only occur in one region at a time), their length was randomly selected among 5, 10 or 100 samples, their amplitude between 50 and 400. This procedure was carried out 47 times, to obtain an independent surrogate dataset for each one of the 47 participants, that will be referred to as the “uncoupled” dataset. The uncoupled dataset was then transformed using the subject-specific leadfield matrix, yielding new surrogate sensor-level timeseries, where each sensor is a weighted sum of all the sources, according to the same leadfield matrix that was used to reconstruct the real data. Noise, correlated as 1/*distance among sensors*, was then added to the sensor-level time series, with a SNR = 4. Then, new source-reconstructed time series were computed for each subject. Based on these new time series, we performed the same procedure to compute the transition matrix as described above. Specifically, we z-scored the time series, thresholded them (threshold z=±3), retrieved the avalanche-specific transition matrices, averaged these within each subject and then across the group, and finally symmetrized the matrix. We then investigated the extent of correlation between the new transition matrix and the structural connectivity matrix. We repeated the entire procedure reported above one hundred times, and show that is unlikely that linear mixing alone can explain the significant association between transition probabilities and structural connectivity (p < 0.001).

### Statistical analysis

The Spearman rank correlation coefficient was used to assess the association between transition probabilities and structural connectivity. A correlation coefficient was computed separately for each individual across all pairs of regions. Transition matrices were symmetrized before this computation. Randomized transition matrices were generated to ensure that associations between transition probabilities and structural connectivity could not be attributed to chance. Avalanches were randomized across time, without changing the order of active regions at each time step. We generated a total of 1000 randomized transition matrices and the Spearman rank correlation coefficient was computed between each randomized matrix and structural connectivity. This yielded a distribution of correlation coefficients under randomization. The proportion of correlation coefficients that were greater than, or equal to, the observed correlation coefficient provided a p-value for the null hypothesis that structure-function coupling was attributable to random transition events.

---

**Supplementary figure 1.**
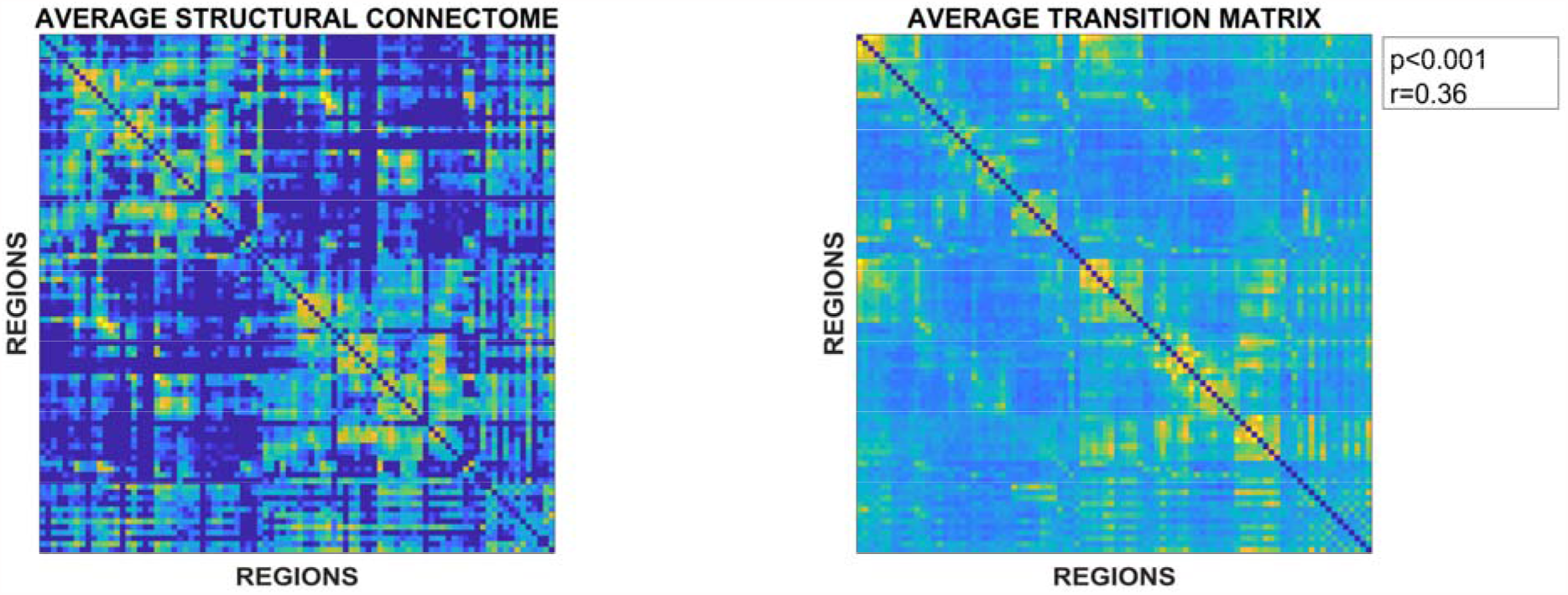
On the left, the average structural matrix. On the right, the average transition matrix.

## Supplementary File 1

**Table.1.** **Correlations between the structural connectome and frequency-specific transition matrices.**

|  | **R** | **p** |
| --- | --- | --- |
| **Delta (0.5 – 4 Hz)** | **0.38** | **2.021e-120** |
| **Theta (4 – 8 Hz)** | **0.35** | **5.34e-100** |
| **Alpha (8 – 13 Hz)** | **0.38** | **1.46e-116** |
| **Beta (13 – 30 Hz)** | **0.38** | **7.30e-122** |
| **Gamma (30 – 48 Hz)** | **0.39** | **1.32e-123** |

### Source data File

The source data file contains the code to generate the transition matrices starting from neuronal avalanches and to compare them to null surrogates.

